# Volumetric studies of phenylalanine in the pre micellar regions of Cetyl trimethyl ammonium bromide (CTAB) in aqueous solution

**DOI:** 10.1101/2023.12.06.570380

**Authors:** Nizamul Haque Ansari

**Affiliations:** Department of Physical Sciences (Chemistry), Sant Baba Bhag Singh University, Jalandhar, Punjab, India-144030

## Abstract

Interactions between amino acid (dl-phenylalanine) and surfactant (cetyltrimethyl ammonium bromide) was investigated using density and density data was utilized to calculate apparent molar volume 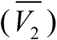, and partial molar volume also known as limiting molar volume 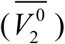 to find out solute-solvent interactions between amino acid and surfactant.

## 1. Introduction

Surfactants are wetting agents that, by lowering a liquid’s surface tension, help spread more easily by reducing the interfacial tension between two liquids. The word “surfactant” was first introduced in 1950 by Antara Products. Among the chemical industry’s most adaptable products are surfactants, which can be found in a wide range of goods, including the motor oils we use in our cars, the prescription medications we take for illnesses, the detergents we use to clean our homes and laundry, the drilling muds used to prospect for petroleum, and the flotation agents used to beneficiate ores [1, 2]. In aquatic environments, surfactants often have a hydrophilic surface and a hydrophobic core [3– 9]. However, phenylalanine is an important amino acid. Amino acids are the building blocks of proteins [10–13]. Foods high in protein contain it. It might enhance learning and memory. It might improve alertness and mood. Information about the thermodynamic and transport characteristics of amino acids in aqueous phase is essential for designing and optimizing the industrial biochemistry processes that are now in use and those that are being proposed. Data on the thermodynamic characteristics of amino acids in aqueous surfactants are uncommon, despite the fact that volumetric measurements of dl-phenylalanine in aqueous solutions are few. Keeping these in mind here we report densities, *ρ*, of dl-phenylalanine (0.00, 0.01, 0.02, 0.03 and 0.04 m) in 0.0001m (mol kg^-1^) aqueous CTAB. The results were interpreted in terms of solute -solute and solute - solvent interaction.

## 2. Results and discussion

### 2.1 Volumetric Study

The experimentally measured values of density (ρ), of 0.015, 0.020, 0.025 and 0.030 m, dl-phenylalanine in 0.0001 m aqueous CTAB at 298.15, 303.15, 308.15, and 313.15K are given in the table 1. The apparent molar volumes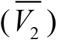, were determined from the solution densities using the following equation:

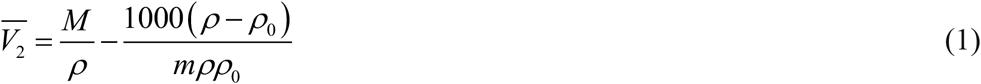

where *M* is the molar mass of the solute, dl-phenylalanine, and m is its molality; ρ_0_ and ρ are the densities of the solvent (aqueous CTAB) and solution, respectively. The calculated values of apparent molar volumes at different molalities of dl-phenylalanine is given in table 2. The limiting apparent molar volumes 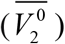 were calculated applying a least-squares technique to the plots of vs. m using the following equation:

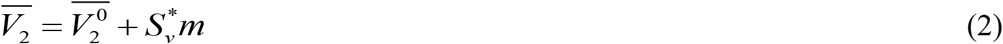

where 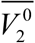 is the partial molar volume at infinite dilution and *S*\*_*v*_ is the experimental slope. The plots 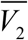 of against the molal concentration (m) were found to be linear with positive 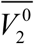. The table 3 shows that 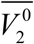 values are positive and increases with temperature, suggesting that solute – solvent interaction increases with the increase of temperature. The transfer molar volumes were calculated by the fallowing relation.

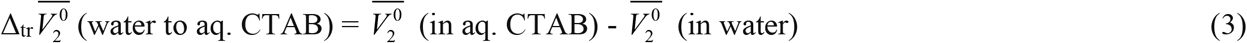

The transfer partial molar volumes are also included in table 3. The table 3 indicates that transfer partial molar volumes are positive with respect to temperature for dl-phenylalanine in 0.0001m aqueous CTAB [14].

**Table 1.** Values of density, ρ, of dl-phenylalanine in 0.0001m aqueous CTAB at different temperatures.

| **m**<br>**(mol kg<sup>-1</sup>)** | **T (K)** |  |  |  |
| --- | --- | --- | --- | --- |
|  | **298.15** | **303.15** | **308.15** | **313.15** |
| **dl-phenylalanine + 0.0001 m aq. CTAB** |  |  |  |  |
| **$\rho$ (kg m<sup>-3</sup>)** |  |  |  |  |
| 0.000 | 1.0006 | 0.9999 | 0.9975 | 0.9958 |
| 0.015 | 1.0014 | 1.0005 | 0.9979 | 0.9961 |
| 0.020 | 1.0021 | 1.0012 | 0.9985 | 0.9966 |
| 0.025 | 1.0030 | 1.0021 | 0.9993 | 0.9973 |
| 0.030 | 1.0041 | 1.0032 | 1.0004 | 0.9983 |

**Table 2.** Values of apparent molar volumes, 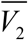, of dl-phenylalanine in 0.0001m aqueous CTAB at different temperatures.

| m<br>(mol kg<sup>-1</sup>) | T ( K) |  |  |  |
| --- | --- | --- | --- | --- |
|  | 298.15 | 303.15 | 308.15 | 313.15 |
| | $\overline{V}_2$ (10<sup>-5</sup> m<sup>3</sup> mol<sup>-1</sup>) | | | |
| dl-phenylalanine + 0.0001 m aq. CTAB |  |  |  |  |
| 0.015 | 11.1730 | 12.5120 | 13.8750 | 14.5670 |
| 0.020 | 9.0046 | 10.0060 | 11.5240 | 12.5450 |
| 0.025 | 6.9040 | 7.7019 | 9.3075 | 10.5220 |
| 0.030 | 4.8395 | 5.5003 | 6.8254 | 8.1644 |

**Table 3.** Values of 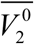 and 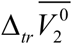 of dl-phenylalanine in 0.0001m aqueous CTAB at different temperatures.

| T/ K |  |  |  |  |
| --- | --- | --- | --- | --- |
| dl-phenylalanine + 0.0001 m aq. CTAB |  |  |  |  |
|  | **298.15** | **303.15** | **308.15** | **313.15** |
| $\overline{V}_2^0 / 10^{-5} \text{ m}^3\text{mol}^{-1}$ | 17.4758 | 19.4327 | 20.89736 | 21.00346 |
| $\overline{V}_{2(aq)}^0 / 10^{-5} \text{ m}^3\text{mol}^{-1}$ | 12.1480<sup>a</sup> | - | - | - |
| $\overline{V}_{2(tr)}^0 / 10^{-5} \text{ m}^3 \text{mol}^{-1}$ | 5.3278 | - | - | - |

a. Millero, F.J; Surdo, A.L; Shin, C., J. Phys. Chem., 1978, 82, 784. Ref. [14].

It has been found [15], that the partial molar volume of a nonelectrolyte is the combination of intrinsic volume of the solute and the volume change due to its interaction with the solvent. Tarasawa et al. [16] pointed out that the intrinsic partial molar volume is considered to be made up of two types of contributions:

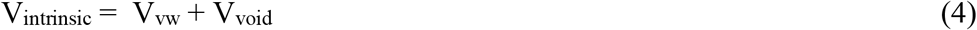

where *V*_vw_ is the volume occupied by the solute due to its van der Waals volume [17], and *V*_void_ is the volume associated with the voids and empty spaces present therein [18]. Shahidi et al. [19] modified the above equation to evaluate the contribution of a solute molecule to its 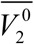 as:

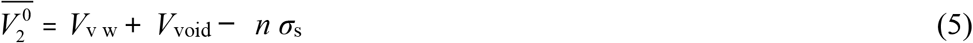

where *σ*_s_is the shrinkage in the volume produced by the interaction of hydrogen – bonding groups present in the solute with water molecules, and n is the potential number of hydrogen – bonding sites in the molecule. Finally, 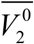 of an amino acid can be viewed as:

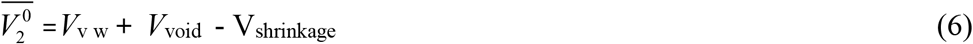

If it is assumed that *V*_vw_ and *V*_void_ are of the same magnitude in water and in aqueous surfactant solutions [20, 21], then the observed changes in 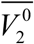 or in 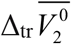 of the amino acids can be explained in terms of changes in the volume of shrinkage in the presence of surfactant molecules in aqueous solutions. As the critical micelle concentration of CTAB [22] is 8.2 × 10^−4^ mol. dm^-3^, the surfactant molecules in this study are present as monomers behaving as electrolyte in the pre – micellar region. If we assume that *V*_v, w_ and *V*_void_are the same in water and CTAB solutions, the positive volume change might arise from a decrease in *σ*_s_in CTAB solution. Because of the presence of CTAB which is cationic in nature, the interaction of the amino acids (dl-phenylalanine) and CTAB is stronger enough, and also the number of hydrogen bonds between amino acid molecules and water molecules increases, thus causing a decrease in *σ*_s_.

